# Distinct composition of plasma virome in HIV-infected subjects on antiretroviral therapy compared to controls

**DOI:** 10.1101/2023.07.21.548740

**Authors:** Tannu Bhagchandani, Mohd. Maksuf Ul Haque, Md Zubbair Malik, Ashwini Kumar Ray, Urvinder Kaur S, Ankita Rai, Anjali Verma, Kamal Kumar Sawlani, Rupesh Chaturvedi, D Himanshu, Ravi Tandon

**Affiliations:** Laboratory of AIDS Research & Immunology, School of Biotechnology, Jawaharlal Nehru University, New Delhi, India; School of Computational and Integrative Sciences, Jawaharlal Nehru University, New Delhi; Host-Pathogen Interaction Laboratory, School of Biotechnology, Jawaharlal Nehru University, New Delhi, India; Laboratory of Metabolic Disorder and Environmental Biotechnology, Department of Environmental Studies, Faculty of Science, University of Delhi, New Delhi, India; Department of Medicine, King George’s Medical University, Lucknow, India; Special Centre for System Medicine, Jawaharlal Nehru University, New Delhi, India

## Abstract

Microbiome study during HIV infection is widely documented with major emphasis on bacteriome, while virome studies are few. The alteration of plasma virome is reported in HIV patients who are either treatment naïve or have a weak immune system. Less is known about the plasma virome in HIV-infected patients on ART with preserved CD4 counts. In our pilot study, viral DNA isolated from plasma was sequenced on Illumina Nextseq500 platform. With the help of VIROMATCH pipeline, we observed that the plasma virome of HIV patients were significantly distinct from controls on the basis of Bray-Curtis dissimilarity index. The species, *Human gammaherpesvirus 4* and families, *Herpesviridae* and *Siphoviridae* were found to be significantly enriched and differentially abundant in HIV patients. Hence, plasma virome is an important component for future study that might influence disease progression among HIV patients during therapy.

## Introduction

Virome is a part of the metagenome that consists of genome of viruses [1]. Viruses are omnipresent that colonize diverse habitats of the planet like air, water, soil, arable lands, etc [1,2]. Humans are continuously exposed to the vast number of viruses in their lifetime, which are a notable fraction of the human virobiota [3,4]. The human viral community is estimated to be about 67.7% of the human body and is composed of eukaryotic DNA and RNA viruses, retroviruses, plant viruses, bacteriophages and newly discovered giant viruses [1,5,6]. The viruses present in humans are not necessarily the ones that cause infections. The composition of virus ranges from acute, latent and chronic infection-causing pathogenic viruses to the ones fairly present in the absence of clinical manifestations [7]. The latter are known as resident or commensal viruses as their presence does not give rise to infections or does not trigger the immune system [4,8].

As of 2020, around 37.7 million people worldwide were living with Human Immunodeficiency Virus (HIV). In India during the same year, 2.3 million people were estimated to be living with HIV, 65% of which had access to antiretroviral therapy (ART). HIV infects and progressively depletes CD4 positive T cells [9]. If not treated, it further progresses to AIDS after passing acute and chronic stages of infection. The commensal virome composition and diversity at various sites changes during HIV infection [10–12]. During HIV infection, microbial translocation due to leaky gut has been identified as a factor contributing to systemic inflammation [13,14]. Besides shifts in bacteria, altered virome composition also contributes to disease progression. Monaco et. al showed an increased abundance of Adenovirus in HIV-infected patients (CD4 counts below 200) whether or not they are on ART [15]. Here, we have analyzed the plasma virome composition of HIV-infected subjects on ART with normal CD4 counts and have compared it with controls.

## Results and Discussion

In this pilot study, blood was withdrawn from six HIV-infected subjects (CD4 count below 200 undergoing ART) and six age-matched uninfected controls to isolate plasma. The viral DNA isolated from plasma was sequenced and the viruses were characterized using VIROMATCH pipeline. The viral reads obtained after analysis belong to single and double-stranded DNA viruses and bacteriophages. The absolute counts obtained were imported into R to calculate relative abundance which was used for downstream analysis. The total number of viral species obtained from our analysis is 275 and their relative abundance in controls and HIV-infected subjects is shown by stacked bar plots (Fig.1a). Further, the highly abundant viral reads found in HIV-infected subjects and controls belong to families - *Anelloviridae, Herpesviridae* and unclassified *Caudovirales* (Fig.1b).

**Fig. 1.**
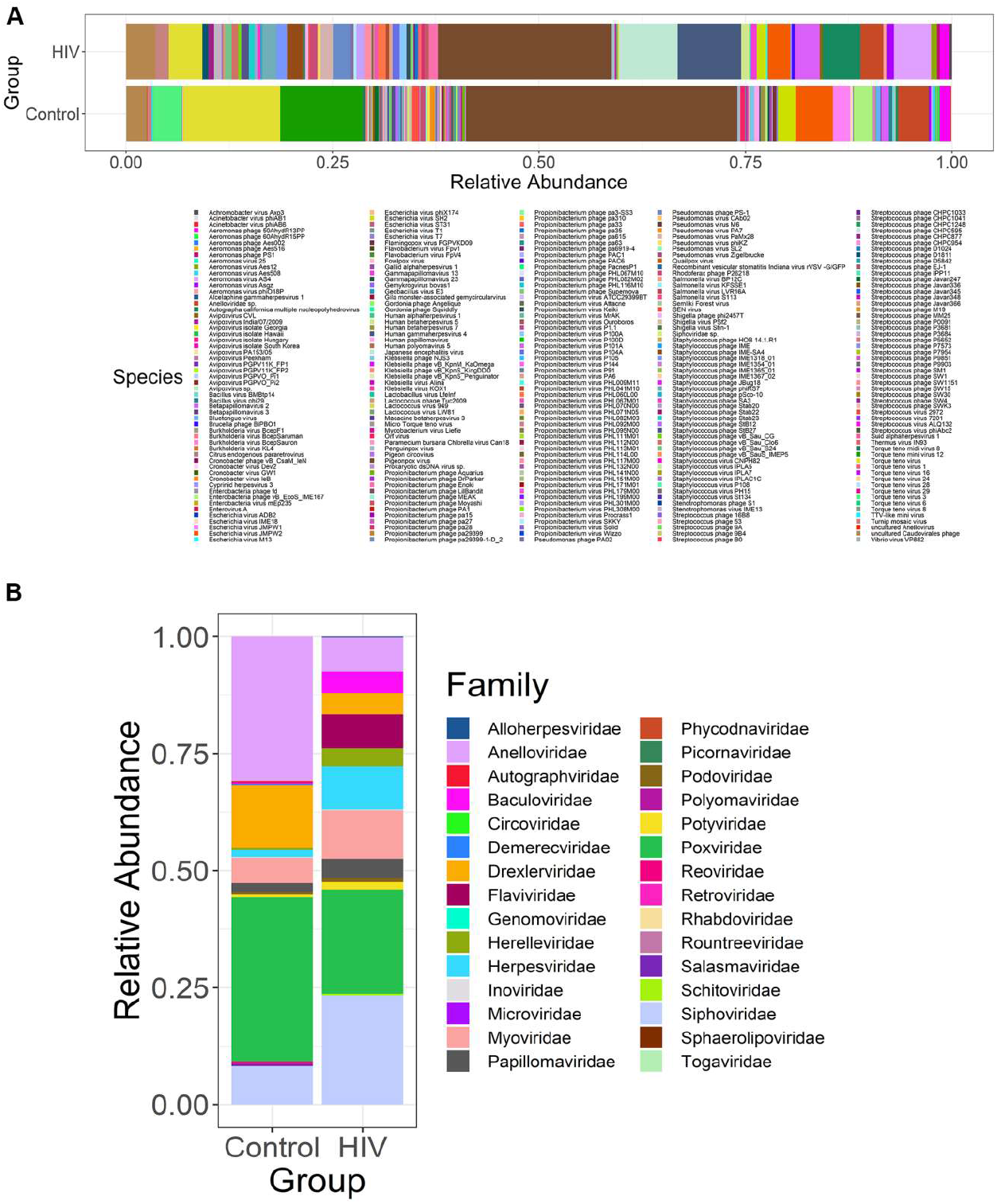
Stacked bar plots showing the relative abundance of viruses at (a) species and (b) family level present in HIV infected patients on ART and uninfected healthy controls.

No difference in alpha diversity, richness and evenness of plasma virome was found between HIV-infected adults and controls. Further, the beta diversity analysis using Principal Component Analysis (PCoA) showed that plasma virome of HIV-infected subjects on ART was significantly different from controls at species and family level (R^2^=0.13944, p=0.012 and R^2^=0.18134, p=0.003, respectively, Fig.2a and b). This suggests that altered plasma virome in HIV-infected adults on ART is either a cause or effect of HIV infection [16].

**Fig. 2.**
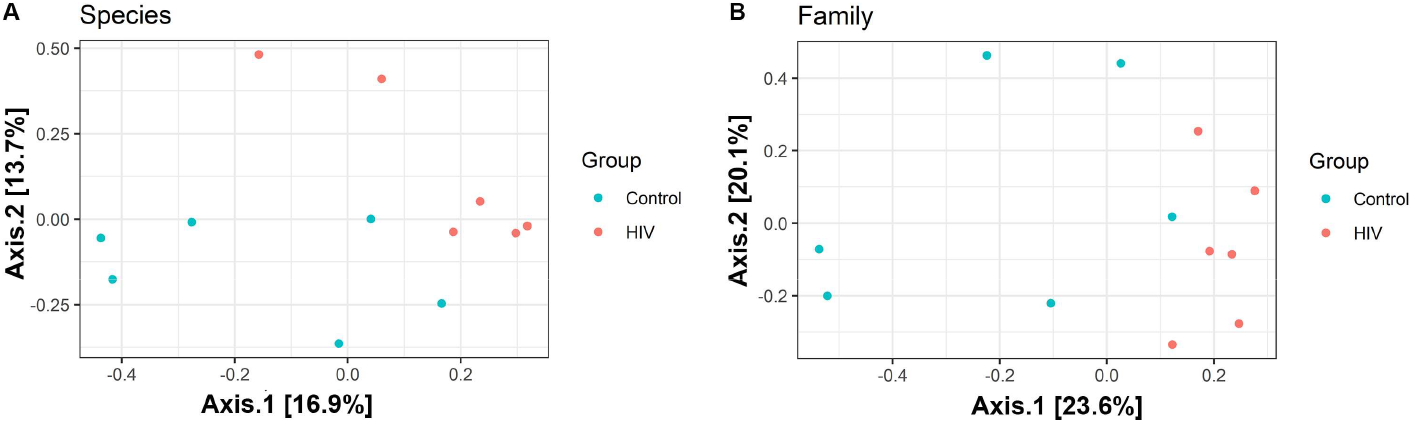
Comparison of beta diversity. PCoA plots representing viral dissimilarities on each PCoA axes between controls and HIV-infected patients on ART at (a) Species (R^2^=0.13944, p=0.012) and (b) Family level (R^2^=0.18134, p=0.003). Statistical signifigance was tested using Adonis test.

In comparison to controls, the relative abundance of the viral species *Human gammaherpesvirus 4* and family *Herepesviridae* and *Siphoviridae* was found to be increased significantly in HIV-infected subjects on ART (p=0.0166, p=0.0368 and p=0.026, respectively) (Fig. 3a, b and c).

**Fig. 3.**
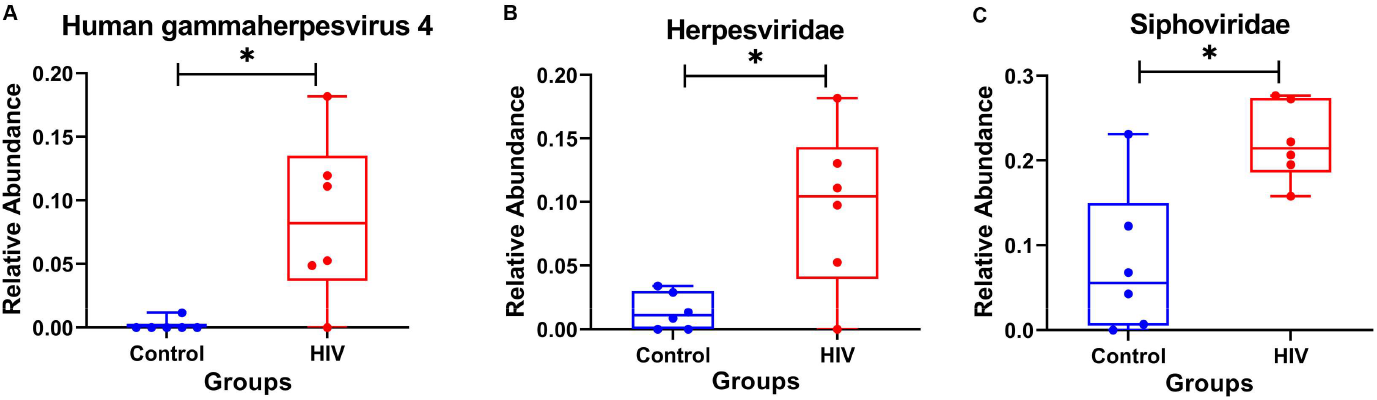
Dot plots representing relative abundance of viruses between groups-controls and HIV-infected patients on ART at (a) Species and (b & c) Family level. Statistical significance was tested using Mann-Whitney test, p<0.05 was considered to be significant.

To identify the viral taxa associated with HIV, we performed LefSe analysis that revealed significant difference in the abundance of 7 taxa between ART-suppressed HIV-infected adults and uninfected healthy controls (LDA score>2.4). LEfSe revealed enrichment of viral species *Human gammaherpesvirus* 4, genus *Lymphocryptovirus*, family *Herpesviridae*, family *Siphoviridae*, order *Herpesvirales*, class *Herviviricete* and phylum *Peploviricota* in HIV-infected subjects undergoing ART (Fig.4a and b). No viruses were found to be enriched in healthy controls at any taxa level.

**Fig. 4.**
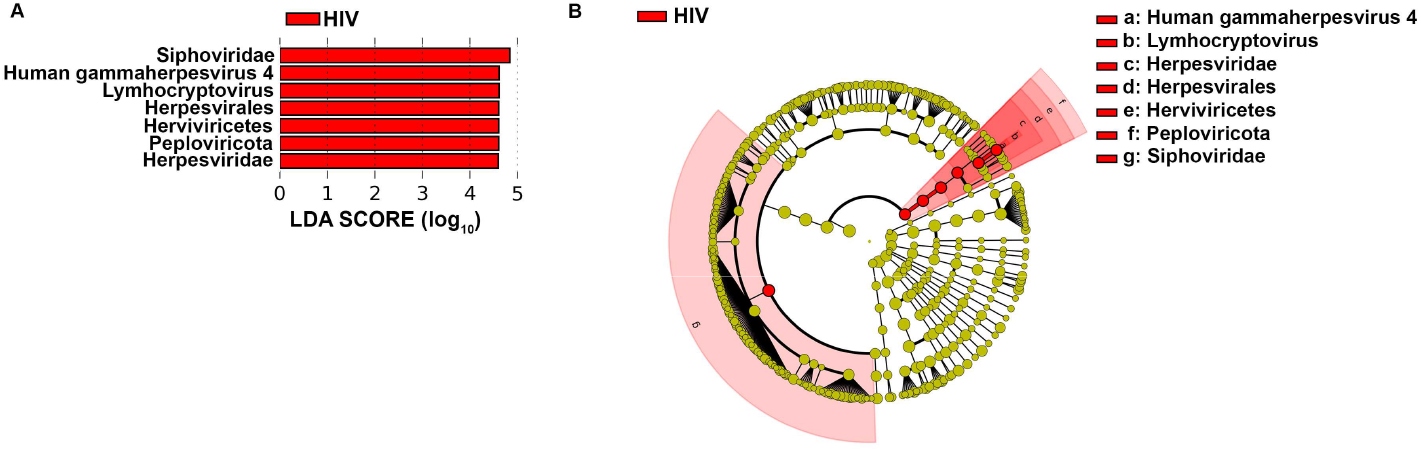
Differentially abundant viruses associated with HIV patients on ART. Linear Discriminant Analysis (LDA) effect size (LEfSe) analysis representing the significantly abundant viral taxa in HIV-infected patients on ART in (a) Histogram representing HIV-infected subjects are shown with positive LDA score. Taxa passing LDA threshold value of >2 are shown. and (b) Cladogram representing the differentially abundant taxa in HIV-infected subjects. Brightness of each dot is proportional to taxon abundance. Red shows viral taxa enriched in HIV-infected adults on ART. Statistical significance was tested using Wilcoxon-Mann-Whitney test, p<0.05 was considered to be significant.

*Human gammaherpesvirus 4*, also known as *Epstein-Barr virus* (EBV) is a herpesvirus belonging to family *Herpesviridae* and is associated with lymphomagenesis. It infects the oropharyngeal route where it mainly targets B cells and epithelial cells [17]. Incidence of EBV is higher in HIV patients [18]. During HIV infection, presence of EBV in PBMC and plasma is regarded as a marker of AIDS defining cancers such as Non-Hodgkin’s Lymphoma (NHL), Burkitt’s Lymphoma (BL) and non-AIDS associated Hodgkin’s lymphoma (HL) [19–21]. Although rates of incidence of AIDS defining cancers have declined since the introduction of ART, NHL and HL still remain a cause of concern in HIV patients on ART [22]. Further, EBV loads have been reported to be higher in ART-suppressed HIV patients. Possible reasons reported for this are partial immune reconstitution during ART and immune activation due to microbial translocation, residual viremia and co-infecting pathogens even during ART [23–26]. Increased abundance of *Human gammaherpesvirus 4* and family *Herpesviridae* in our study are thus in line with previous findings.

In conclusion, the plasma virome profiles in HIV infected patients on ART were found to be distinct from controls. The difference could be due to *Human gammaherpesvirus 4* as evidenced by its differential abundance in HIV infected patients on ART. Limitations of our study are the small sample size and less explored nature of virome due to the small and incomplete virus database.

## Acknowledgements

Authors would like to acknowledge DST-Fund for Improvement of S&T Infrastructure (FIST)(DST File NO:SR/FST/ LS1-653) for supporting Flow cytometry and sorter facility. Authors are thankful to Indian Council of Medical Research (ICMR), Government of India (61/6/2020-IMM/BMS); Department of Biotechnology (DBT), Government of India. RT acknowledges UGC-Faculty Recharge Programme (UGC-FRP).

